# aHISplex: an imputation based method for eye, hair and skin colour prediction from low coverage ancient DNA

**DOI:** 10.1101/2023.11.02.565295

**Authors:** Zoltán Maróti, Emil Nyerki, Endre Neparaczki, Tibor Török, Gergely István Varga, Tibor Kalmár

## Abstract

The prediction of externally visible traits (eye, hair and skin colours) from DNA can provide valuable information for both contemporary and ancient human populations. The validated HIrisPlex-S method is the primary tool in forensics for phenotyping modern samples. The HIrisPlex-S multiplex PCR assay can handle trace DNA from modern samples, but the analysis of degraded, low coverage ancient DNA (aDNA) presents additional challenges. Genotype imputation has recently proven successful in effectively filling in missing information in aDNA sequences. To assess the feasibility of this approach, we evaluated how key factors, such as genome coverage, minor allele frequency, extent of post mortem damage, and the population origin of the test individual influence the efficiency of imputing HIrisPlex-S markers and predicting phenotypes. We used high coverage sequence data from ancient remains for the evaluation. Our results demonstrate that even with genome coverages as low as 0.1-0.5x, the proposed workflow is capable of predicting phenotypes from degraded ancient (or forensic) WGS data with good accuracy. To aid the archaeogenetics community, we have developed a user-friendly, easily deployable imputation-based framework that includes the new bioinformatics tools and the pre-made reference data sets required for the whole analysis.

## Background

The prediction of externally visible human traits from DNA has implications in forensics [1–3] but it can also provide valuable information for the analysis of archaeological remains. The most comprehensive and forensically validated HIrisPlex-S system currently uses 41 autosomal markers to simultaneously predict eye, hair, and skin colour [4–6]. While the HIrisPlex-S multiplex PCR assay can cope with trace amounts of DNA in forensic samples, the analysis of ancient DNA (aDNA) presents further challenges. DNA from archaeological remains is prone to post mortem damage (PMD), leading to DNA degradation over time. Typically, aDNA consists of very short DNA fragments, with an average length of 40-60 base pairs, and it undergoes frequent C>T and G>A nucleotide transitions, primarily near the ends of these fragments. These characteristics, including the small average DNA fragment size, potential contamination with high molecular weight modern DNA and frequent nucleotide changes, render PCR-based methods (including the HIrisPlex-S multiplex PCR assay) unfeasible on ancient samples. Consequently, majority of the population genetic analyses of aDNA are based on shotgun WGS or hybridisation-based targeted methods.

To date, a substantial number of WGS sequences have been generated from ancient samples to facilitate the genetic characterization of ancient populations. However, due to the persisting challenges in sequencing degraded aDNA, there are currently only a handful of high-coverage WGS sequences available from aDNA, while the majority of aDNA samples typically have low genome coverage (0.1x-2x). Although the current HIrisPlex-S system can handle partial information, its primary reliance on diploid genotype data results in increased uncertainty in phenotype predictions and a higher likelihood of producing invalid results when dealing with missing or low-confidence diploid genotypes. Due to the low genome coverage of aDNA samples, a significant portion of HIrisPlex-S markers typically lacks sequence information. Although sparse pseudo-haploid genotypes from aDNA can be used in a number of population genetic approaches like PCA, admixture and qpAdm, this data is not suitable for phenotype assessment with the HIrisPlex-S system.

With the increasing number of low-coverage WGS sequences, new tools and approaches are emerging to address the challenges of analysing incomplete data. One of the latest approaches is to impute the missing diploid genotypes/haplotypes from partially genotyped data. This approach relies on the observation that closely linked markers tend to be inherited together as haplotypes from parents to offspring. Hence, a catalogue of such common haplotypes and the partial information on sequential stretches of markers can be used to predict the most likely diploid state of markers in the low coverage sample. Several algorithms and tools have been developed to balance speed and precision in addressing the large computational requirements of the imputation [7–9]. Available literature indicates that imputation precision depends on multiple factors, including the fraction of missing and genotyped markers, the minor allele frequency of the imputed marker, the density of genotyped markers, the reference data set, the genome structure/population origin of the imputed sample, and the choice of algorithm/tool [8-10].

State-of-the-art imputation tools can achieve high genotyping precision for common variants with high minor allele frequency (MAF), even when dealing with partial information corresponding to 0.5x genome coverage data. Since most genetic markers of eye, hair, and skin colours consist of common alleles with high MAF, imputation has the potential to predict the diploid genotype state of these markers even from low-coverage aDNA. Given that the three predicted phenotypic traits result from the combination of several trait-defining markers, we hypothesised that a few imputation errors would not significantly impact the prediction of the most likely phenotype. This suggests that the approach can be feasible for phenotyping ancient individuals from aDNA.

We developed a user-friendly, readily deployable imputation-based framework containing the new bioinformatics tools and the pre-made reference data sets required for the analysis. In our proposed workflow we selected the last version of GLIMPSE tool (GLIMPSE2, version 2.0.0) for imputation with the gold standard phased 1KG reference data, as it promises very fast and accurate imputation compared to other tools [10, 11]. Using our framework, we assessed the effect of major influencing factors (the genome coverage, minor allele frequency, extent of PMD, and the population origin of the test individual) on the imputation efficiency and phenotype prediction using high coverage experimental data from ancient remains.

## Results

We expanded the genome windows around each of the 41 HIrisPlex-S marker coordinates by 2.5 million base pairs in both the 5’ and 3’ directions. Imputation was performed on the 1.7 million common variants within the resulting 11 genome segments, covering a total of around 70 megabases, as described in the methods section. To assess imputation accuracy and the robustness of imputation-based genotyping, we selected 31 high-coverage (>10x genome coverage) ancient WGS shotgun sequences from the Allen Ancient Data resource (AADR, version 54.1) [12, 13]. The sample IDs, the associated annotations from the AADR database, and the global population structure based on the FastNGSAdmix tool [14] of the selected samples are found in Supplementary Table S1. From the high-coverage WGS data, we obtained the most likely genotypes of the 31 samples for all 41 HIrisPlex-S markers, except for a few positions where they had 5x or lower genome coverages (Supplementary Table S2). We utilized the diploid genotypes from the high-coverage data as the ground truth for assessing concordance and genotyping errors in imputation.

The original high-coverage data was downscaled to 2x, 1x, 0.5, and 0.1x genome coverages in 10 parallel replicates, each with different random seeds. Next, we conducted 10 parallel imputations for each downscaled dataset, resulting in a total of 100 parallel imputed sets of genotypes for each sample at each genome coverage. We filtered the imputed genotypes of each 41 HIrisPlex-S marker from all imputed variants and compared them with the ground truth to calculate the fraction of imputation errors. As imputation is a probabilistic method, we also tested the correspondence between the imputed genotypes from the original high coverage data and the ground truth. Thus, we also imputed the diploid genotypes from the original high-coverage samples in 100 parallel runs (Supplementary Table S3).

We distinguished between two types of genotyping errors based on their potential impact on phenotype prediction. The ‘total error’ encompassed all cases where the imputed diploid genotype differed from the ground truth. The ‘opposite error’ refers to instances where homozygous opposite alleles appeared in the imputed sequence compared to the truth (HOM REF instead of HOM ALT or vice versa), as the second category would more severely affect the phenotype prediction probabilities considering either dominant-recessive, or quantitative inheritance.

### Effect of genome coverage on imputation accuracy

According to our results, the mean imputation error rate was most severely affected by the genome coverage (Figure 1.), as this factor influences the number/density of genotyped markers used to predict the most likely haplotype configurations and diploid genotypes at all loci.

**Figure 1.**
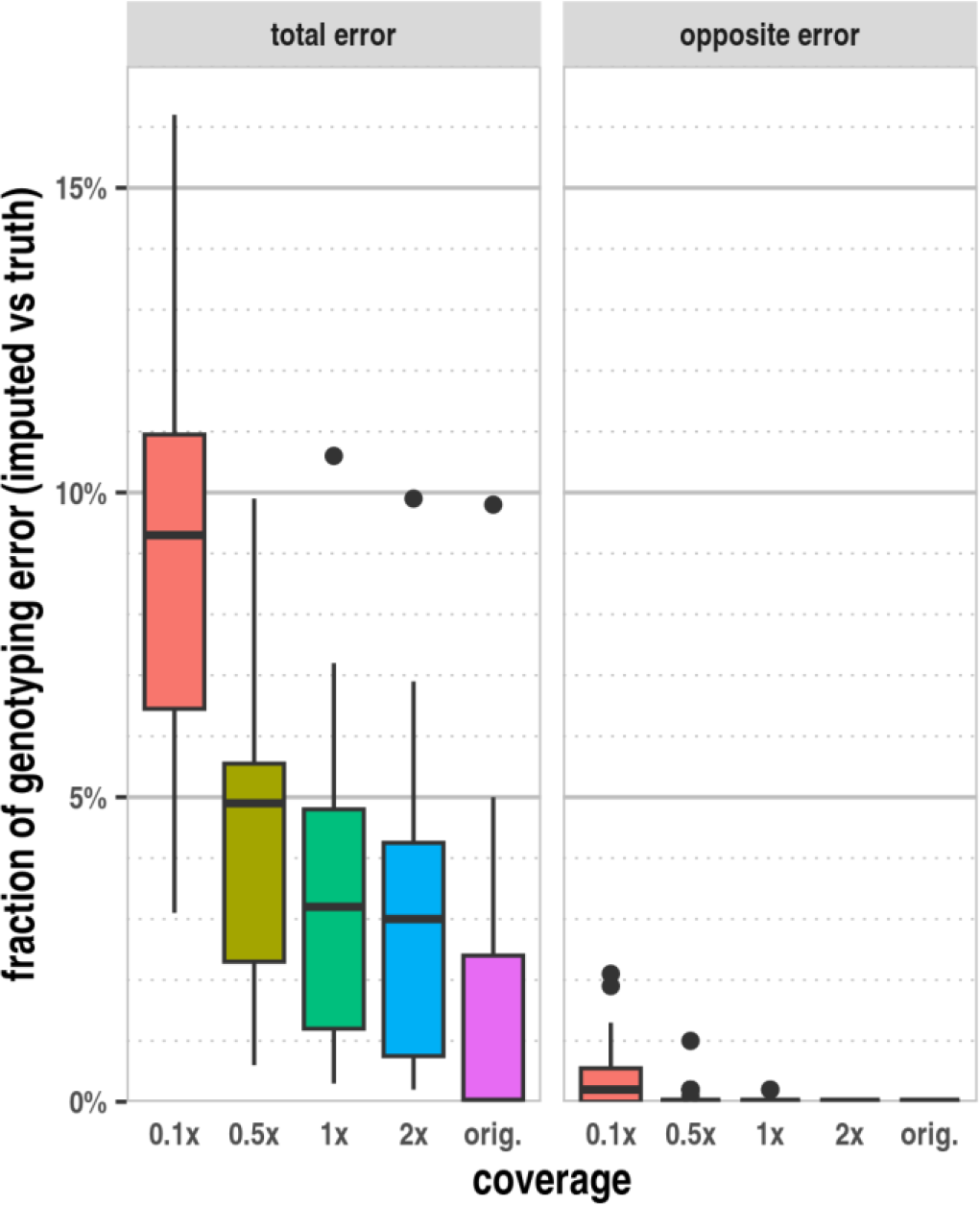
The effect of genome coverage on the mean imputation error based on the analysis of 100 parallel imputations of 31 ancient samples. In the “total error” category any imputed genotype that did not match the expected ground truth was considered. In the “opposite error” category we only considered cases were homozygous opposite alleles appeared in the imputed sequence compared to the truth.

The accuracy of the imputed diploid genotypes is noteworthy. The median concordance of imputed diploid genotypes with the truth is approximately 95% at 0.5x and approximately 91% at 0.1x genome coverage for the analysed samples. At higher genome coverages the accuracy increases up to 97%. Nevertheless, even in the original high coverage data a very low frequency of imputation errors can be observed, likely at variants with skewed allelic balance. The rate of opposite genotype errors is negligible at or above 0.5x genome coverages, and even at 0.1x genome coverage, only a few samples had more than 1% of this type of error. At 1x genome coverage and below, most markers are usually represented by a single read, with only one allele randomly sampled. At 0.5x genome coverage, statistically half of the markers are covered, nevertheless the overall diploid genotyping accuracy remains 95% or higher. This high accuracy demonstrates the feasibility of imputing diploid genotypes for the HIrisPlex-S markers from ancient, low-coverage shotgun WGS data.

### The effect of sample age, population origin and genotyping errors on imputation accuracy

We also calculated the error rates in different sample subgroups to assess other factors that can impact imputation accuracy. These factors included sample age (which affects post mortem aDNA damage), the use of Uracil*-*DNA glycosylase (UDG) treatment (as it drastically reduces the amount of transition errors in the sequences), and the sample’s source population(s) (as the reference data and the genome structure of the test population could be largely different). Unfortunately, the number of available high-coverage ancient samples does not allow us to create homogenous groups based on all potentially influencing factors. Therefore, we created contrasting subsets of samples (containing at least three samples) that differed in one or more influencing factors. We then calculated the mean genotyping error rates for these subsets (Table 1).

**Table 1:** Mean imputation error rates observed in the different subsets of samples at different genome coverages based on 100 parallel imputations. UDG – Uracil*-*DNA glycosylase; EUR – European origin; AFR – African origin; BP – before present.

|  | N | mean imputation genotyping error fraction (%) |  |  |  |  |  |  |  |  |  |
| --- | --- | --- | --- | --- | --- | --- | --- | --- | --- | --- | --- |
|  |  | total error |  |  |  |  | opposite error |  |  |  |  |
|  |  | 0.1x | 0.5x | 1x | 2x | orig. | 0.1x | 0.5x | 1x | 2x | orig. |
| ALL | 31 | 9.0% | 4.4% | 3.4% | 2.9% | 1.4% | 0.4% | 0.1% | 0.0% | 0.0% | 0.0% |
| no UDG (ALL) | 15 | 9.2% | 4.4% | 3.6% | 3.1% | 1.9% | 0.4% | 0.1% | 0.0% | 0.0% | 0.0% |
| UDG (ALL) | 16 | 8.7% | 4.4% | 3.3% | 2.8% | 0.9% | 0.5% | 0.0% | 0.0% | 0.0% | 0.0% |
| EUR (noUDG) | 3 | 11.4% | 6.6% | 5.4% | 3.8% | 1.6% | 0.4% | 0.0% | 0.0% | 0.0% | 0.0% |
| EUR (UDG) | 9 | 9.5% | 4.1% | 2.5% | 2.1% | 0.5% | 0.6% | 0.0% | 0.0% | 0.0% | 0.0% |
| AFR (UDG) | 3 | 7.6% | 6.0% | 6.0% | 5.4% | 3.3% | 0.3% | 0.1% | 0.1% | 0.0% | 0.0% |
| 30000-41400 BP | 3 | 10.4% | 4.9% | 4.0% | 3.3% | 0.8% | 0.8% | 0.2% | 0.1% | 0.0% | 0.0% |
| 8240-9615 BP | 4 | 8.6% | 2.8% | 2.4% | 1.4% | 0.9% | 0.1% | 0.0% | 0.0% | 0.0% | 0.0% |
| 163-1170 BP | 5 | 6.5% | 4.7% | 4.3% | 3.6% | 2.0% | 0.3% | 0.1% | 0.0% | 0.0% | 0.0% |
| EUR (ALL) | 12 | 9.9% | 4.7% | 3.2% | 2.6% | 0.8% | 0.5% | 0.0% | 0.0% | 0.0% | 0.0% |
| AFR (ALL) | 4 | 8.0% | 5.6% | 5.5% | 5.0% | 4.9% | 0.3% | 0.1% | 0.1% | 0.0% | 0.0% |
| OTHER (ALL) | 15 | 8.5% | 3.9% | 3.0% | 2.6% | 0.9% | 0.3% | 0.1% | 0.0% | 0.0% | 0.0% |

The detailed data of the imputation genotype error rates for each individual sample at the different genome coverages are shown in Supplementary Table S4. Overall, the mean error rates in the selected subgroups exhibit only minor differences, underscoring that the primary factor influencing imputation errors is the genome coverage. It is noteworthy that samples of African origin (AFR) exhibit higher imputation error at higher genome coverages, even when using the original high coverage data. This observation strongly suggests the presence of reference errors, stemming from the fact that the greater haplotype diversity among AFR populations are not adequately represented in the 1KG reference panel. It appears that UDG treatment slightly decreases the imputation error rate, as evidenced by the difference between UDG treated and non-UDG treated individuals of European origin (EUR). This effect is likely associated with the higher level of PMD in samples without UDG treatment, which leads to an increased rate of random C>T and A>G transitions in ancient DNA sequences. We also observe a slight increase in the mean imputation error among older dated samples. As sample age in general correlates with the PMD level in aDNA, it is likely that this increase in imputation error rates is attributable to the age factor. Importantly, the opposite genotype error rate remains negligible at 0.5x genome coverage levels or higher, even in the oldest samples at 0.1x genome coverage, the opposite error rate did not exceed 0.8%.

### The imputation accuracy of the individual HIrisPlex-S markers

To evaluate whether certain markers exhibit higher imputation error rates due to low MAF, insufficient linked marker context or poorly represented haplotypes in the reference data, we also calculated the mean genotyping error rate for each individual marker (Supplementary Table S5). Overall, most of the 41 HIrisPlex-S markers, including those with lower MAF, had good imputation accuracy. Nevertheless, we noticed that certain markers had markedly worse imputation accuracy in specific samples even at high genome coverages.

In order to distinguish between high and low imputation error rates, a threshold value needs to be defined. We established this threshold based on the currently available imputation accuracy for African genomes. In the literature the highest (>= 15%) imputation error rates are observed in African samples, even at higher genome coverages, and these are clearly associated with reference bias. Therefore, in our analysis, we considered the mean imputation error at high genome coverages (original, 2x and 1x) exceeding 15% as “high” error rates, and those below this threshold as “low” error rates. Applying this criteria, we identified 55 sample/marker combination that had significantly elevated imputation error rates across the original, and all lower coverage imputations (Supplementary Table S6). Out of the total of 1271 sample/marker combinations (31 genomes with 41 markers) this small subset of combinations accounted for ∼47% of all imputation errors, as indicated by the number of imputation errors in the second row of the high error rate category in Table 2.

**Table 2.** Summary of high and low error rate sample / marker combinations. The criteria of >= 15% mean imputation error frequency observed in the high genome coverage (original, 2x, 1x) imputations was used to distinguish between high/low imputation error rate combinations.

|  | high error rate sample/rsID combinations<br>(N=55)* |  |  |  |  | low error rate sample/rsID combinations<br>(N=1217) |  |  |  |  |
| --- | --- | --- | --- | --- | --- | --- | --- | --- | --- | --- |
| genome coverage | original | 2x | 1x | 0.5x | 0.1x | original | 2x | 1x | 0.5x | 0.1x |
| number of imputed genotypes | 5500 | 5500 | 5500 | 5500 | 5500 | 121600 | 121600 | 121600 | 121600 | 121600 |
| number of imputation errors | 1748 | 2470 | 2735 | 2818 | 2793 | 0 | 1219 | 1618 | 2814 | 8610 |
| % of imputation errors (error/no. of imputations*100) | 31.78% | 44.91% | 49.73% | 51.24% | 50.78% | 0.00% | 1.00% | 1.33% | 2.31% | 7.08% |
| fraction of total error in the given category (high/low error rate vs total error) | 100.00% | 66.96% | 62.83% | 50.04% | 24.49% | 0.00% | 33.04% | 37.17% | 49.96% | 75.51% |

Imputation determines the best likelihood of all genotype data within a large chunk of analyzed genome region of the test individual that fits to the best matching diploid allele combination inferred from the reference data. Discrepancies between linked markers in the sample and linkage information in the reference data can lead to inevitable ambiguities or incorrect predictions of the diploid genotype. Our data also suggests that within this smaller subset of combinations the primary cause of the elevated error rate is likely reference bias. Most prominently, imputation error occurs even in the original high coverage data. Furthermore, within the high-error category, the overall frequency of imputation errors remains significantly elevated at every level of genome coverage. This is in contrast to the low-error category of sample/marker combinations, where errors tend to increase as genome coverage decreases (3^rd^ row in Table 2).

### Imputation analysis of the rs312262906 red hair marker

The low minor allele frequency of the imputed marker is known to lead to lower concordance. While most HIrisPlex-S markers have high minor allele frequencies, the less common red hair trait is associated with low minor allele frequency genetic variations [15]. The HIrisPlex-S system contains seven rare (MAF<0.002) markers to predict the red hair trait, rs312262906, rs11547464, rs1805008, rs1805006, rs1805009, rs201326893, and rs1110400. The red hair marker with lowest global MAF of 0.00078 is the rs312262906 variant, which, unlike all other HIrisPlex-S markers is not a single nucleotide polymorphism (SNP). Instead it is a one base pair duplication (NC_000016.10:g.89919344dup) resulting in an additional adenosine nucleotide insertion in the sequence. The typically lower genotyping accuracy of small insertions or deletions compared to unique SNPs, coupled with the extremely low MAF could potentially have a significantly adverse impact on the genotyping and imputation accuracy of this marker.

As we had shown, the genotyping error rates of the individual markers showed similar figures for the rare markers. However, since none of the analysed ancient samples carried any red hair markers, this only shows that the false positive imputations of these low MAF markers were not excessive. Therefore, we also aimed to investigate the true positive recall of a known red-hair individual carrying this sequence variation. Due to the very low global MAF of this marker, only 5 unrelated individuals have this variation in the GRCh38 aligned 1KG phase III data set. Consequently, we selected data from a single high-resolution modern sample (NA10830) from the 1KG phase III data that carry a heterozygous mutation for this marker as our test individual.

Since GLIMPSE2 is a reference-based imputation method, we created a custom GRCh38 reference data set that excluded the data from NA10830. We also verified that none of its relatives was included, as it would invalidate the results. We followed the same experimental setup as described for the ancient samples: the original ∼30x NA10830 data were downscaled to 2x, 1x, 0.5x, and 0.1x genome coverages in 10 parallel runs using different random seeds, and imputation was performed in 10 parallels for each downscaled dataset.

According to our analysis, the imputation of genotypes for this individual was 100% accurate down to 0.5x genome coverage for all markers, including rs312262906. Even at 0.1x genome coverage, 80 out of 100 parallel runs resulted in the correct heterozygous dupA for rs312262906 without additional genotyping errors of the other markers. Our analysis revealed that, in the case of this high-quality modern sequence imputation was highly deterministic. For the same downscaled data, the inferred genotypes from parallel imputations were consistently identical (Supplementary Table S7). This suggests that in the downscaled 0.1x coverage BAM files with false negative imputation for this particular marker, all supporting reads containing the linked markers (and the rs312262906 marker itself) that are crucial for imputing accurate genotypes, were absent.

### Phenotype classification accuracy of the imputation-based data

The HIrisPlex-S system calculates the probabilities of each phenotypic trait using a model that was cross-validated on a large dataset comprising both genotypic and phenotypic information. The web tool translates the genotype information of markers into a set of p-values [16], which can subsequently be used to classify each phenotypic trait. Due to the varying number of phenotype associated markers and the HIrisPlex-S classification scheme, the number of phenotype categories differ for the three phenotypic traits. The classification of the eye phenotype (blue, brown, black) is straightforward, as the highest phenotype probability (p blue eye, p brown eye, p black eye) indicates the most likely eye colour. In contrast, the classification of hair colour (red, blonde, dark-blonde/blonde, dark-blonde/brown, brown, dark-brown/brown, dark-brown/black, black) and especially the skin phenotype (very pale, very pale/darker, pale/lighter, pale, pale/darker, intermediate/lighter, intermediate, intermediate/darker, dark/lighter, dark, dark/dark-black, dark-black) is more complex. It relies on a greater number of phenotype-associated markers and involves complex heuristic rules based on the calculated p-values, as described in the HIrisPlex-S web tool manual [17]. To simplify and streamline phenotyping, we have incorporated a tool in our software package (*classifHISplex*), which classifies the three phenotypic traits based on these rules and the p-value output file from the HIrisPlex-S web tool.

To evaluate the impact of imputation errors on phenotype classification, we conducted a comparison between classifications based on the genotypes from high coverage data (considered as ground truth) and those based on the 100 parallel imputed genotypes at various genome coverages for each ancient sample (Supplementary Table S8). In general, a smaller number of phenotype-associated markers tend to result in statistically fewer imputation errors but potentially have a greater effect on the predicted phenotype, whereas a larger number of phenotype-associated markers tend to lead to statistically more imputation errors but with less drastic changes in the phenotype classification. For both eye and hair phenotypes, the majority of parallel imputations yielded an exact matching phenotype classification with the ground truth, even at 0.1x genome coverages (Figure 2).

**Figure 2.**
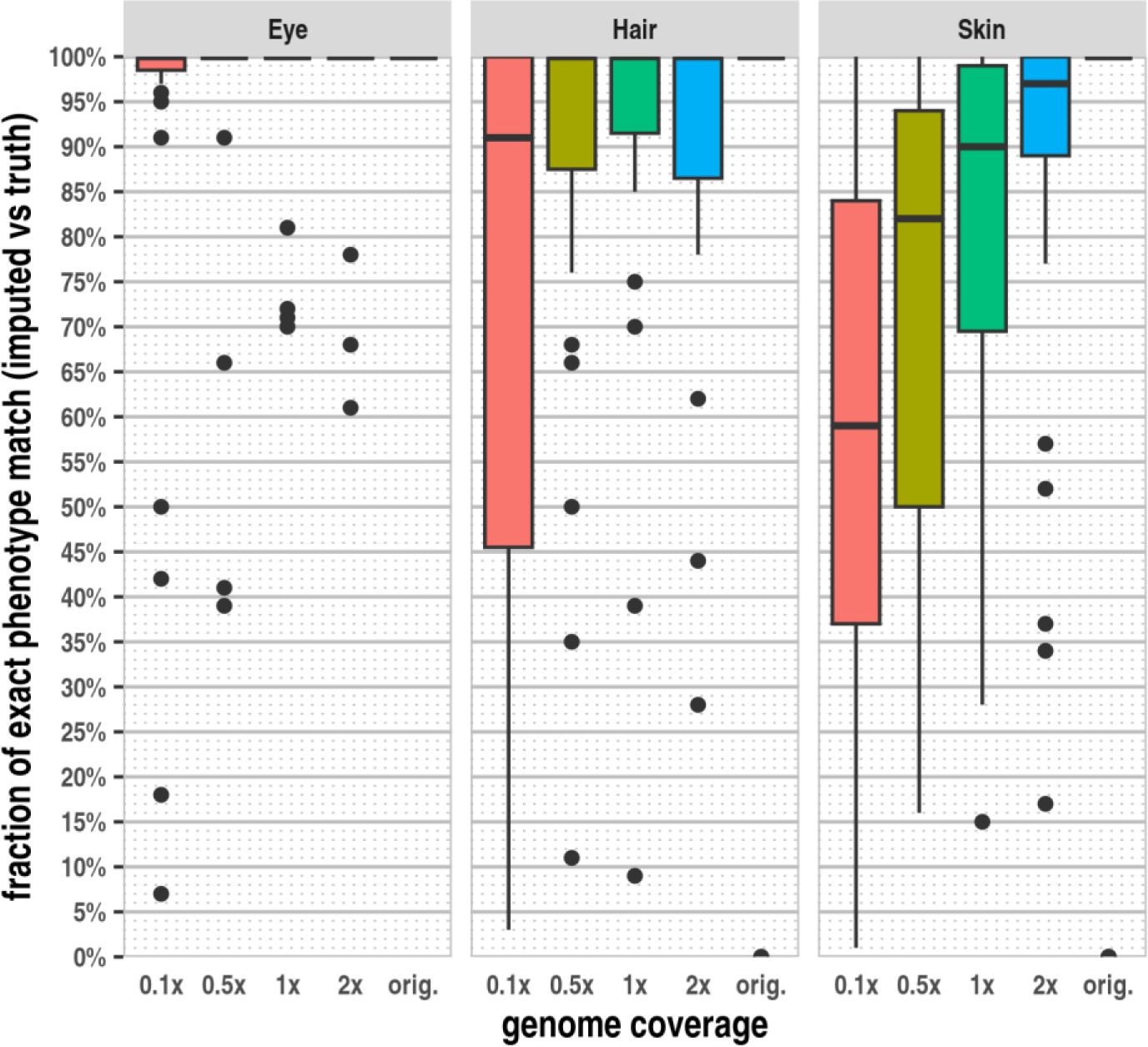
The fraction of exact matching phenotypes for the three phenotypic traits. The phenotypes based on the genotypes of 41 HIrisPlex-S markers from the original high-coverage data were used as the ground truth for each sample. We compared the true phenotypes with those based on the 100 parallel imputated genotypes and calculated the ratio of exact matching phenotypes for each sample at each genome coverage.

The fraction of exact matching skin phenotypes was lower, particularly at lower genome coverages and higher imputation error rates. Although the fraction of exact phenotype classifications is lower in case of the skin phenotype even at higher imputation error rates, a spectrum of very similar phenotypes, compared to the truth, is indicated. Meanwhile, the number of significantly different phenotypic classifications remains relatively low. Our data suggests that this is primarily due to the finer granularity of classification in the case of the skin phenotype. Our results demonstrate that the observed rate of imputation errors, even at very low genome coverages, has a relatively minor impact on the phenotype probability values and the classification of intermediate phenotypes. However, for recessive traits, such as blue eye, blond hair or very pale skin, an imputation error, even in one of the underlying homozygous recessive state markers, leads to darker phenotype and a more severe misclassification. While we had only one individual with dark blonde hair and no individuals with very pale skin among the high genome coverage ancient samples, five out of the 31 samples were classified as having blue eyes. In these samples, the correct blue eye phenotype was predominantly indicated in the majority of parallel imputations at genome coverage 1x or higher. For two samples (Loschbour.DG and VK1_noUDG.SG), the majority of imputations yielded the correct blue eye phenotype (91% and 100% respectively) even at 0.5x genome coverage. In contrast, for the other three samples (vbj004_noUDG.SG, SF12.SG and SZ45.SG) at 0.5x genome coverage, the blue eye phenotype was misclassified as brown with a high probability (61% and 59%, 34% respectively, Supplementary Table S8). As expected, the misclassification rate of the blue eye phenotype was even higher at 0.1x genome coverage. When investigating the eye phenotype associated markers, our data revealed that the higher-than-expected rate of imputation error in latter samples was attributed to sample (population) specific reference bias of the rs12913832 marker (Supplementary Table S6). The three particular samples (vbj004_noUDG.SG /Sweden Gotland Vasterbjers Pitted WareBattleAxe/, SF12.SG /Sweden Mesolithic/, SZ45.SG /Hungary Langobard outlier/) likely share similar marker combinations in this locus, which are poorly represented in the reference dataset. In these particular samples the homozygous A/A minor allele is imputed as heterozygous A/C in approximately 30% of the cases, even at 1-2x genome coverages, signifying reference bias.

## Discussion

The vast majority of shotgun aDNA sequences consist of low genome coverage data, typically ranging from 0.1x to 2x coverage. Due to the low genome coverage, not all genome positions are covered with reads, leading to sparse genotype data. Furthermore, the low genome coverage does not permit true diploid genotyping, as in most positions only one allele is (randomly) sequenced. The sparse, pseudo-haploid genotypes are not suitable for DNA-based phenotyping because the diploid state of the phenotype-defining markers is required to predict the exact phenotype with high confidence. Consequently, despite the abundance of available aDNA sequences, because of this limitation, we have phenotypic information only for a handful of ancient samples.

Imputation of the most likely haplotypes from sparse pseudo-haploid data promises to uncover the diploid genotypes with high confidence. While a very recent manuscript has shown that imputed diploid genotypes have high overall concordance even from noisy, low-quality, and low genome coverage ancient samples [18], there is currently no data available to assess whether this approach can reliably be used for the evaluation of complex phenotypic traits in ancient samples. Another challenge is that the entire workflow, from aligned NGS data to phenotypic classification involves numerous tools, data preparation and data shaping, with no single straightforward tool available to perform the entire analysis. Consequently, our goal was to create a user-friendly tool to facilitate the imputation-based workflow and to evaluate the effect of the major influencing factors on the proposed workflow. Accordingly, we investigated how the genome coverage, genotyping errors, sample age and the overall genome structure (population origin) of the sample influence imputation errors and the accuracy of phenotype prediction.

There are different approaches used for imputation [7–9]; however, the majority of imputation tools infer the common haplotypes from phased, fully typed diploid references. While some approaches can assess common haplotypes ‘on the fly’ using a vast number of jointly analysed samples, these tools typically require very large sample sets and higher CPU resources. Unfortunately, such large, high-quality sample sets are not yet available for ancient samples. Therefore, despite their limitations, reference-based imputation methods appear to be the only feasible option for analysing ancient samples. The imputation accuracy of reference panel-based methods inevitably depends on the reference used, as the representation of common haplotype configurations (series of linked markers) serve as a model to impute the most likely haplotypes of the test sample. The assumption that our gold standard reference data set represents most of the likely haplotypes in our test data clearly imposes limitations on the approach. Consequently, imputation can potentially yield false haplotypes/genotypes that exist in our reference but not in the test data. This can occur, for example, in the case of young markers that did not exist yet at the given date in the ancient sample, or when the genome structure of the test individual is not (properly) represented in the reference dataset.

Our results have confirmed that the most significant portion of the imputation errors could be attributed to genome coverage, as it directly impacts the quantity, quality and ploidy of genotype information used to infer the most probable haplotypes. In general, the mean genotyping error rate in the analysed ancient genomes was less than 5% even at genome coverages as low as 0.5x. Additionally, we demonstrated that the opposite genotype error was nearly non-existent at 0.5x or higher genome coverages, only becoming detectable at 0.1x genome coverage. Our data also confirms that even when dealing with partial genotype data, where only random single alleles are available due to low genome coverage, diploid genotypes can be imputed with high accuracy (Figure 1).

According to our results there were only minor differences in the mean imputation error rates among different populations (Table 1). Previous literature has shown that the 1KG phase3 reference data set is less representative for populations of African origin, leading to a higher imputation error rate [9, 10, 18]. The mean imputation error rate at 0.5x genome coverage, considering all variants in whole genome sequences, is expected to be 15 % or higher for genomes of African origin [18]. However, in the case of the analysed HIrisPlex-S markers the observed mean genotyping error was only ∼5.6% at this genome coverage in AFR samples (Table 1). It is known that many markers associated with pigmentation are under selection pressure in Africa [19, 20], leading to less genetic diversity and likely better representation of haplotypes at these loci. This effect could contribute to the observed low imputation error rate of these markers in the majority of the tested African origin individuals. A notable difference was that AFR individuals had a considerably high imputation error rate even at 1x or 2x genome coverages, indicating reference bias and falsely imputed markers due to improper haplotype representation for a few HIrisplex-S markers in the used 1KG phase3 reference (Supplementary Table S6).

When comparing the overall imputation error rates among different HIrisPlex-S markers, we observed that some of the markers exhibited considerably higher error rate across all genome coverages (Supplementary Table S5). Our analysis revealed that a portion of the imputation error is not random. Specifically, we identified 55 high error rate sample/marker combinations that collectively accounted for approximately ∼47% of all imputation errors. Our results indicate that only specific samples had very high imputation errors at specific markers, even at the original, 2x and 1x genome coverages, while the rest of the samples had very low error rates for these markers (Supplementary Table S6, Table 2). When the reference is not representative for the haplotype combination of specific samples, we can expect a high deterministic imputation error. Only about 5% of all sample/rsID combinations have high imputation error that can be attributed to reference error. Our data indicates that, for the majority of imputed genotypes, the imputation error frequency caused by partial genotype information is consistently low, even when the genome coverage is as low as 0.5x.

Markers of the HIrisPlex S system have moderate to high minor allele frequencies, except for those associated with red hair. Previous literature has indicated that imputation concordance tends to be lower for rare variants. Therefore, we conducted an accuracy test for the red hair variant (rs312262906), which has an exceptionally low minor allele frequency (MAF). In our study, using a high-quality genome of modern European origin, we observed a very high true positive recall rate for rs312262906. At genome coverage of 0.5x or higher, genotype imputation achieved 100% accuracy. Even at 0.1x genome coverage, this marker was correctly imputed in 80% of the parallel runs, while for the other HIrisPlex S markers, diploid genotypes were inferred with 100% accuracy at 0.1x coverage (Supplementary Table S7).

The probabilities of each phenotype, including 3 eye colours, 7 hair and 11 skin tones, are determined based on the set of underlying genetic markers and HIrisPlex-S prediction models. Since these probabilities are contingent upon the combination of multiple markers, it is expected that in most cases, the observed 1-2 imputation errors out of the 41 HIrisPlex-S markers do not significantly impact the phenotype probabilities and the derived phenotypic classifications. This is particularly true for intermediate phenotypes, where the p-values may shift slightly either direction, but this typically does not result in substantially different phenotype probabilities and classifications. In principle, recessive traits such as blue eye, blonde hair, and very pale skin are more susceptible to severe misclassification, as imputation error in any of the underlying markers can result in a darker (dominant) phenotype being assigned. Our results show that majority of imputation replicates still correctly classified blue eye at 1x genome coverage or higher. However, at 0.5x genome coverage three out of 5 samples with blue eyes experienced higher misclassification rate compared to the correct classification (Supplementary Table S8). Our findings revealed in these three ancient northern European samples, the primary cause of imputation errors and subsequent misclassification was the reference bias of the rs12913832 marker. While imputation errors equally impact the markers associated with darker shades, the heuristic applied in the phenotype classification scheme, which gives precedence to the darker tones, mitigates this issue, thus the classification of individuals with very dark pigmentation is less severely affected.

Ultimately, the 31 samples were properly classified for the three phenotypic traits in the majority of parallel imputations, even at 0.1x genome coverages. The skin colour classification showed slightly larger phenotypic variability, but the predicted skin tones showed only minor differences compared to the truth (Supplementary Table S8). The higher incidence of observed minor differences in the indicated skin tone can be likely attributed to the more finely graded skin phenotype classification, which relies on a larger number of skin tone-associated genetic markers and involves more complex heuristic decision rules for deriving probabilities.

The HIrisPlex-S system requires true diploid genotypes for each of its 41 markers to predict phenotypes. It is noteworthy that at 0.5x genome coverage, statistically, only half of the markers have pseudo-haploid genotype information available, and at 0.1x genome coverage, only 4-5 markers are covered by a single read. Despite this significant amount of missing information, the genotype imputations and the phenotype predictions were surprisingly accurate. Our analysis underscores the strength of imputation and demonstrates that even at these extremely low genome coverages, the applied method yields comparable results to phenotype predictions derived from the original high-coverage data.

## Conclusion

In summary, we have developed an easily deployable, user-friendly, imputation-based workflow that incorporates the validated HIrisPlex-S system for phenotyping ancient samples. This proposed workflow is practical for analysing ancient whole-genome sequencing (WGS) data, even at low genome coverages of 0.1x to 0.5x, with the expectation of accurate classification in the majority of cases. Our results suggest that modern sequences with no genotyping errors and a closely matching reference lead to improved imputation accuracy, even when dealing with extremely low coverage. Currently, only a handful of ancient samples have been phenotyped due to the absence of suitable tools. Our workflow and the tools we’ve developed now enable the analysis of approximately 1500-2000 publicly accessible ancient WGS datasets that possess sufficient coverage to predict the phenotypes of past populations. Since a significant portion of the challenges stem from reference bias, it is anticipated that as reference genomes become enriched with high-quality data, this workflow could achieve even higher concordance rates. Furthermore, as our understanding of genotype-phenotype associations continues to expand, the proposed workflow can be readily extended to accommodate new phenotypic markers. Additionally, aside from ancient samples, this framework can be employed for the analysis of degraded forensic samples, where the HIrisPlex-S multiple PCR-based method is hindered by short DNA fragments.

## Methods

### Used data sets

We selected 31 publicly available, high coverage ancient samples from the Allen Ancient Dataset (AADR, version V54.1) [12, 13]. Specifically, we chose samples that underwent shotgun whole-genome sequencing (WGS) and had genome coverage exceeding 10x. The annotations of the 31 selected samples, including information about their origin, and sample type can be found in Supplementary Table S1. The selected samples include both UDG and non-UDG treated samples, originates from various geolocations (including samples with EUR and AFR ancestry), and span a wide range of dates from 41400BP to 1920CE. In all of our downstream analyses, we used the high coverage alignment files (BAM) available in public repositories referenced in the original manuscripts.

### Simulation of low coverage data

We computed the total average genome coverage for each high-coverage ancient genome using mosdepth [21]. To simulate low genome coverage we used ***samtools*** [22] with the appropriate “***view -- subsample 0*.*FRAC --subsample-seed INT***” options to downscale the original high coverage BAM files. From each high coverage BAM file, we generated ten parallel downscaled 2x, 1x, 0.5x and 0.1x genome coverage data sets, each with different (seed 1-10) random seed.

### Preparation of reference marker sets

We prepared GRCh37, hg19, GRCh38, hg38 reference data sets to analyse the 41 HIrisPlex-S markers. Using the appropriate genome coordinates, we created a BED file that contains the marker positions for each reference data. Using “***bedtools slop***” [23], we extended the genome windows around each marker by 2.5M base pairs in both the 5’ and 3’ directions. Then we merged the overlapping genome windows using “***bedtools merge***”. If the resulting genome window was smaller than 5 million base pairs in size (in case of markers near telomeres or centromeres), we extended the genome window in the opposite direction. As a result, each of the resulting genome windows were at least 5 million base pairs, ensuring an ample number of flanking markers around the 41 HIrisPlex-S markers. We used GLIMPSE2_chunk with the “*–recursive*” option to create a list of genome regions suitable for imputation. We used the biallelic SNPs of the appropriate 1KG Phase 3 reference data. As the GLIMPSE2 framework only imputes genotypes at positions that exist in the reference data, we manually included the genotypes of the single biallelic dupA variant for red hair (rs312262906) to the reference. The GLIMPSE2_split_reference tool was used to generate the binary reference files required to impute the variants in the 11 genome regions containing the 41 HIrisPlex-S markers.

### Detailed description of the aHISplex tool and workflow

We created an easily deployable package (https://github.com/zmaroti/aHISplex) that contains all the required tools, customised reference data and translation tables for the different reference genomes to run the whole analysis workflow from aligned BAM file(s) to phenotype classification. The workflow consists of three steps.

- The first step is to run the GLIMPSE2 based imputation on a single BAM file or list of BAM files and filter/translate the imputed genotypes of the HIrisPlex-S markers resulting in a HIrisPlex-S web tool compatible data file. The tool also saves all the underlying raw analysis files and the log files of GLIMPSE2 phasing and ligation.
- In the second step the ‘*HISplex41_upload*.*csv*’ output file generated in the first step (conforming with the required HIrisPlex-S upload data file format) has to be uploaded and analysed at the official https://hirisplex.erasmusmc.nl/ website using the HIrisPlex-S batch upload phenotyping web service.
- In the third step the downloaded results file can be processed by the ***classifHISplex*** tool (included in our software package) to evaluate the resulting phenotype probabilities and classify each analysed sample for the three phenotypic traits based on the rules described in the HIrisPlex-S manual [17].

The tools of the aHISplex software package are written in golang and a shell script is included as a glue to call and run the public (GLIMPSE2_phase, GLIMPSE2_ligate, bcftools and optionally GNU parallel tool) and the included **t*ransToHISplex*** tool with the appropriate parameters. The whole analysis workflow (except the web tool part) can be performed with two commands providing the BAM file(s) and the output of the web tool. The github page contains a detailed README for the dependencies, installation and usage of the package.

### Assessment of aHISplex performance on the HIrisPlex-S system

We downscaled each sample for each genome coverage in ten parallel runs with different (1-10) random seeds. Thus, the available genotype information (how many reads and at which ratio the alleles are represented) for the imputed markers was different in each downscaled data set. Furthermore, even from the same information, based on the allele counts and haplotype probabilities in the test sample and the most probable haplotype from the reference the predicted genotypes, multiple different solutions may exist (with different probabilities) leading to different imputed genotypes. Thus, we performed ten different imputations using each ten parallel downscaled data for each genome coverage and sample, resulting in 10 (imputation parallels) * 10 (downscale parallels) * 5 (different genome coverages) analysis for each of the 31 analysed sample.

We genotyped the high coverage original BAM files based on the allele pileup (***samtools mpileup***) at genome positions of the 41 HIrisPlex-S markers. Using the genotypes, the strand and test allele information of the HIrisPlex-S markers, we calculated the allele counts for each marker and sample (Supplementary Table S2) and used this information as ground truth for the assessment of the imputation genotype concordance.

For the 10x10 parallel imputation-based genotype predictions we calculated the genotype concordance with the ground truth. We counted total differences including all genotypes that were imputed differently, and we also counted the gross error situation, where an HOM ALT truth was imputed as HOM REF (or the opposite). Furthermore, since imputation can result in different genotypes from the same input data, we also calculated the count/frequency of minor alleles (imputation variability).

Using the 10x10x5 imputation based allele counts of the 41 HIrisPlex-S markers for each sample, we run the phenotype probability calculation on the HIrisPlex web server to obtain the probability scores of the predicted hair, eye and skin colours (Supplementary Table S3).

We used the included ***classifHISplex*** tool to classify the phenotype probabilities to eye, hair and skin colour shades based on the rules defined in the “HIRISPLEX-S, HIRISPLEX & IRISPLEX Eye, Hair and Skin colour DNA Phenotyping web tool USER MANUAL” [17].

## Ethics approval and consent to participate

Not applicable.

## Consent for publication

Not applicable.

## Availability of data and materials

We used publicly available data for the validation of the method. Our software with potential future updates is available at the GitHub repository (https://github.com/zmaroti/aHISplex).

## Author contributions

Conceptualization, methodology, software, formal analysis Z.M. Resources E. NY., G.I.V., E. N. Interpretation of results Z.M., T.K. and T.T. Writing original draft Z.M. and T.K. and T.T. All authors took part revising the results and contributed to the final manuscript.

